# Resolution of Reprogramming Transition States by Single Cell RNA-Sequencing

**DOI:** 10.1101/182535

**Authors:** Lin Guo, Xiaoshan Wang, Mingwei Gao, Lihui Lin, Junqi Kuang, Yuanbang Mai, Fang Wu, He Liu, Jiaqi Yang, Shilong Chu, Hong Song, Yujian Liu, Jiadong Liu, Jinyong Wang, Guangjin Pan, Andrew P. Hutchins, Jing Liu, Jiekai Chen, Duanqing Pei

## Abstract

The Yamanaka factors convert mouse embryonic fibroblasts (MEFs) into induced pluripotent stem cells (iPSCs) through a highly heterogeneous process. Here we profile single cells undergoing an optimized 7-day reprogramming process and show that cells start reprogramming relatively in sync, but diverge into two branches around day 2. The first branch of cells expressing *Cd34/Fxyd5/Psca* become nonpluripotent. The second one contains cells that are first *Oct4*+, then *Dppa5a*+ and pluripotent. We show that IFN-γ blocks this late transition. Our results reveal the heterogeneous nature of somatic cell reprogramming, identify *Dppa5a* as a marker for pluripotent and innate immunity as a potential barrier for reprogramming.

**One Sentence Summary:** Single cell RNA sequencing reveals a continuum of cell fates from somatic to pluripotent and *Dppa5a* as a marker for chimera-competent iPSCs.

## Introduction

Reprogramming of somatic cells to a pluripotent state represents a breakthrough for both regenerative medicine and biology (*1-5*). For regenerative medicine, reprogramming enables the generation of patient-specific functional cells that can help cure diseases such as Parkinson’s disease and spinal cord injury (*6-11*). For basic biology, reprogramming has provided valuable insight into how cell fate and cell fate transitions are regulated (*12, 13*). By analyzing the cellular and molecular processes associated with reprogramming, it is clear that the process follows various states with distinct molecular signatures (*14-19*), that have been mapped by comprehensive transcriptomic, proteomic and epigenetic studies (*16, 17, 20-25*), generally only on bulk populations of cells. The analysis of bulk samples suggests MEFs are induced to a pluripotent state passing through several distinct biological processes, such as an early mesenchymal-epithelial transition, autophagy, histone and DNA demethylations (*12, 19, 26-28*).

Despite progresses made through bulk analyses as outlined above, very little is known at the single cell level. The averaging of populations of cells tends to mask infrequent occurrences during reprogramming, thus obscure very rare essential cellular transitions, or overemphasize irrelevant biological processes not required for reprogramming. The fact that only a small fraction of the starting cells eventually become pluripotent demands a vigorous reappraisal of principles and mechanisms established by bulk analysis at single cell resolution (*12, 29*). On the other hand, single cell analysis may be able to help us uncover rare but important mechanisms that have evaded so far (*30-32*). In this report, we take advantage of an efficient and accelerated reprogramming system that converts MEFs to chimera competent iPSCs within seven days. By analyzing this reprogramming process at the single-cell resolution by RNA sequencing, we show a cell fate continuum bifurcating as early as day 2 towards either successful iPSC generation or alternative fates. Along this continuum, we have identified additional barriers at the very late phase of reprogramming, and also critical factors, particularly *Dppa5a*, that appear to govern the final transition to pluripotency. Further analysis at single-cell resolution will provide deeper insights into cell fate decisions that may be generally applicable to both biology and diseases.

### Chimera competent pluripotent cells from OSK infected MEFs in 7 days

We take advantage of our previously described iCD1-OSK (OCT4, SOX2, KLF4) reprogramming system with MEFs carrying the OG2 (*Oct4*-GFP) reporter. This system results in around 10% of cells becoming iPSC colonies, and GFP+ cells begin to appear as early as day 3 (*33*). However, as *Oct4*-GFP is not a marker for complete reprogramming, we decided to evaluate if any of the GFP+ cells are chimera competent. To this end, we infected MEFs with retrovirus expressing *Oct4, Sox2 and Klf4* (OSK), and *Oct4*-GFP positive cells were isolated by FACS sorting, or picked-up at different days (See methods). These isolated GFP+ cells were injected into mouse blastocysts directly without culturing (Fig. 1A). This assay allows us to conclude that we can generate chimera competent cells around day 7 or 8 (Fig. 1B, Table S1 and S2). The resulting chimeras indicated that day 7 GFP+ cells are chimera competent as 11.9% (7 out of 59) of live pups were chimeras (Fig. 1, B and C).

**Fig 1.**
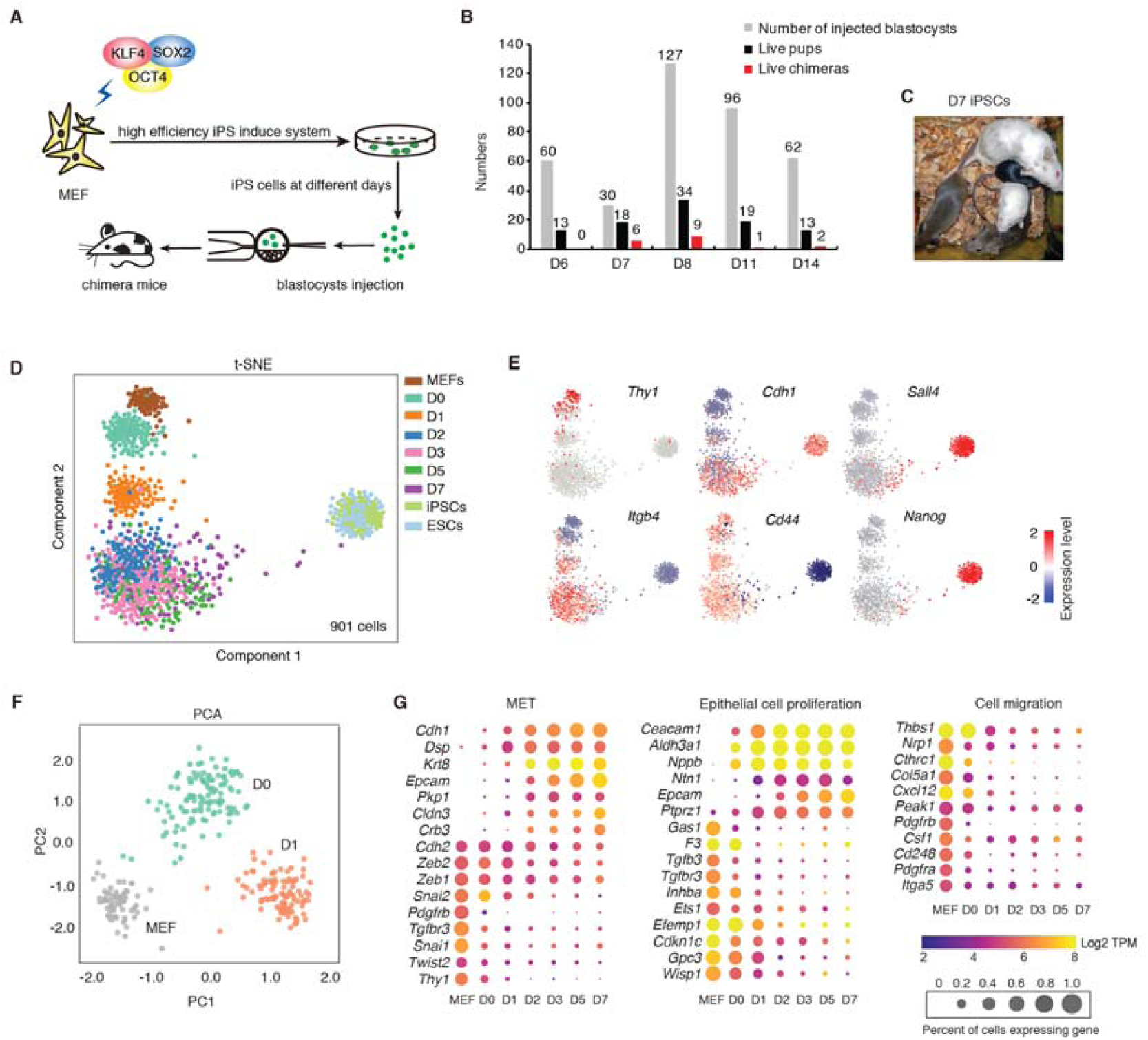
Highly efficient reprogramming system with a chemical defined medium and single cell RNA sequencing recapitulates the known features of reprogramming. **(A)** Schematic diagram of OCT4/SOX2/KLF4 induced reprogramming and assay for chimera competency. iPSCs were isolated by FACS sorting or directly picked at different days, and injected to mouse embryos to identify the pluripotency. **(B)** Chimera competent iPSCs are first detected at post-infection day 7 (D7) of a reprogramming time course. More details see Table S1. **(C)** Chimeric mouse with germline transmission ability were derived from iPS colonies at D7. **(D)** t-SNE projection of all the single cells sequenced, and reveals cell fate transition during reprogramming. Colors indicate different cell time points/conditions. **(E)** tSNE plot from Fig. 1D colored based on expression level of marker genes. **(F)** PCA projection based on the top 3000 high variance genes among MEF, D0 and D1 time points. **(G)** The expression pattern of representative high PC loading genes in Fig. 1F. The size of dots indicates expression percentage in the corresponding time point/conditions (TPM > 2 is considered as expressed), and the colors indicate mean expression level in the cells expressing the respective gene.

### Single-cell RNA sequencing

After ascertaining chimera-competent iPSCs in this reprogramming system, we collected single cells with the C1 platform during reprogramming. In total, we isolated 1045 single cells at D0, D1, D2, D3, D5, D7, plus MEFs, iPSCs and ESCs (fig. S1A), and 901 of them passed quality control. We detected a median of 6889 genes and a minimum of at least 2982 genes in each cell (fig. S1, B to D). Pearson correlation between the mean expression for all cells at each time point to the bulk RNA-seq samples showed good correlation (R>0.8) (fig. S1E). We also estimated the infection efficiency of each exogenous transgenes by using reads from the RNA-seq that align cross the CDS and LTR (fig. S2A). We found that around 90% of the MEFs expressed all three exogenous transgenes, and the correlation between the expression level of the three transgenes was high (fig. S2, B and C). Projection of the RNA-seq expression into a t-SNE space helped visualize the transitions as the cells reprogram (Fig. 1D). Compared to the tight clustering of single cells from MEFs and ESCs, the reprogramming cells are more scattered (Fig. 1D). When key genes were mapped onto the t-SNE, it is clear that cells transition through distinct phases from *Thy1*+ to *Nanog*+ states in 7 days (Fig. 1E).

### Reprogramming begin at near synchrony

Based on the projection in Figure 1D, it is apparent that cells at D0 and D1 are tightly clustered. Indeed, this is consistent with PCA analysis based on the top 3000 high variance genes among MEF, D0 and D1 (Fig. 1F). Plotting the gene contributions to PC1 and PC2 indicates that the left side of the PC1 loading represents mesenchymal-related genes, consistent with our earlier finding that reprogramming starts with a mesenchymal to epithelial transition (MET) (*19*) (fig. S2D). *Thy1* was reported before as a marker for cells resisting to reprogramming (*16*), but it was almost completely silenced in our system at D1 (Fig. 1, E and G, and fig. S2E). Interestingly, the right side of the PC1 loading represents genes critical for metabolism (fig. S2D), suggesting that cells at D1 are leaving the mesenchymal state and reaching a metabolically active intermediate state. We further confirmed the well-defined MET process, which should serve as a validation assay for our single cell data (Fig. 1G and fig. S2F). Furthermore, consistent with previous report that cell surface markers such as CD73, CD104/CD49d are expressed by intermediate reprogramming prone cells (*34*), we show that *Nt5e* (coding for CD73) and *Itgb4* (coding for CD104) are upregulated in most of cells by day 2, and *Itga4* (coding for CD49d) at day 1 (fig. S3A). The upregulation of *Itgb4* is positively correlated to an early epithelial program, as measured by the co-expression of the epithelial cadherin Cdh1 (fig. S3, B and C). Additionally, although *Itg4b* is only expressed briefly at day 1, we show that CD104+ cells are biased towards successful reprogramming (fig. S3, D and E).

### Reprogramming cells bifurcate early into reprogramming vs non-reprogramming cells

In order to understand the fate trajectories of cells undergoing reprogramming, we combined pathway and gene set over dispersion analysis (PAGODA) (*35*) with diffusion mapping analysis (*36*). As shown in Figure 2A, two fate branches emerged. By loading the 519 differentially expressed genes between them on a heatmap, we show that one branch is likely to represent reprogramming fate (R, downward) as these cells express mostly pluripotent genes such as *Tet2*, *Sall4*, and *Dppa5a,* while the second branch represent the non-reprogramming (NR, upward) fate in which cells express *Fxyd5, Psca* an *Cd34* (Fig. 2A, 2B). When we mapped specific genes to the diffusion map, it is clear that while *Cdh1* appears to be present in both NR and R cells, *Cd34*, *Fxyd5* and *Psca* with the NR, *Sall4* and *Dppa5a* with R (Fig. 2C). We also analyzed the cellular trajectories using monocle2 (*37, 38*), which also generated two branches (NR’ and R’, respectively) that are highly similar to the NR/R branches described above (fig. S4, A to E). One explanation for the differences between R and NR fates could be the differential expression of reprogramming factors from viral-delivered transgenes. However, we found no correlation between the expression levels of exogenous transgenes and R/NR fates (fig. S5, A and B). In fact, NR cells express similar levels of OSK as R cells (Fig. S5A), suggesting that the bifurcation into R and NR is not due to the absence of OSK infection in the NR cells.

**Fig 2.**
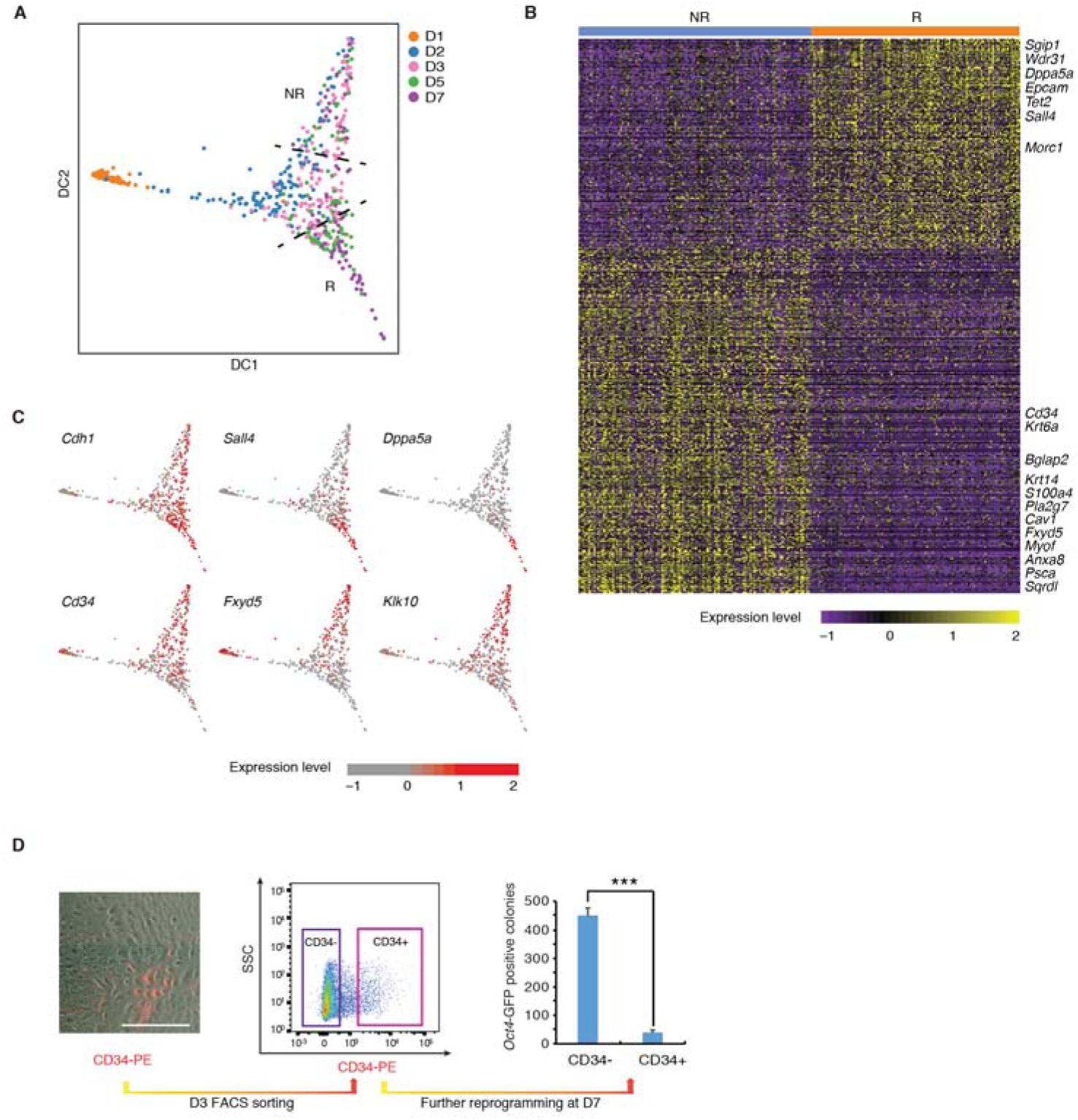
A divergent non-reprogramming trajectory in OSK induced reprogramming. **(A)** Trajectory reconstruction of D1-D7 cells reveals two fates of MEF reprogramming. Each dot corresponds to a single cell, and each color represents the sampling time point. The black dash line above defines a non-reprogramming branch (NR) and the dash line below defines the reprogramming branch (R) respectively. 555 cells from D1 to D7 were used as input and the transcriptional heterogeneity was enriched by PAGODA. DC, diffusion components. (**B).** Heatmap shows the significant differentially expressed genes between the NR and R branches. Significance was determined at an adjusted p value < 0.001 from DESeq2. **(C)** Expression of marker genes representing the NR and R branches shown using the same layout in Fig. 2A. Each point is a single cell and is colored based on the intensity of expression. **(D)** CD34 positive and negative cells were separated by FACS at post-infection D3 (middle), and replanted for further reprogramming. After replanting, *Oct4-*GFP positive colonies were analyzed at post-infection D7 (right), n = 3. Scale bar, 500 µm, ***p < 0.001.

Given the fact that genes such as *Psca* and CD34 clearly mark the NR and NR’ identified in both methods (Fig. 2B, Fig. S4B and S4E), we decided to take advantage of the fact that CD34 being a well-known cell surface marker to show that CD34+ cells do not overlap with *Oct4*-GFP positive cells at D7 (fig. S5C). To see if CD34 can denote the bifurcation at early time point, we sorted D3 cells into CD34+ and CD34- population and replanted them for further reprogramming (Fig. 2D and fig. S5D). As shown in Figure 2D, when scored as *Oct4-*GFP positive colonies at D7, we show that the CD34+ cells are 10 times less efficient than the CD34- ones to give rise to GFP+ iPSCs, indicating that CD34 marks cells for the NR branch. Flow cytometry analysis of further reprogramming cells at D7 also showed that most of the CD34+ population at D3 remained CD34+ cells, and hardly acquired *Oct4*-GFP, whilst CD34-cells rarely gained CD34 expression, indicating that CD34 cells are committed to an NR fate (fig. S5D). We then randomly picked Oct4-GFP+ and GFP- colonies at day 9 and performed RNA-seq on them. We found that the *Oct4*-GFP negative colonies express gene set from non-pluripotency lineages, indicating that they have reached a stable alternative fate during reprogramming process (fig. S5E).

### Approaching pluripotency

A significant portion of the R cells also fail to reach pluripotency based on the scattered nature of the D7 cells in Figure 1D. To further deconstruct the fate of these cells, we revisited Figure 1D and identified 5 single cells that appear to have committed to pluripotency (Fig 1E, 3A and fig. S6A). These 5 cells are very closely related to the the ESC group, and expressing a series of pluripotent genes such as *Dppa5a*, *Tdgf1* and *Utf1* (fig. S6B). Indeed, when we clustered the D7 cells into 6 groups with an unsupervised SNN-Cliq method (*39*) (Fig. 3B), we could independently identify these 5 cells shown in Figure 3A from the rest. The five pluripotent cell (PC) candidates show a strong similarity to ESCs in a co-correlation heatmap analyzed by SC3 (REF) (Fig. 3C), suggesting that these cells may be the ones capable of contributing to chimeras as demonstrated in Figure 1B. Indeed, the expression patterns of critical pluripotency genes in D5, D7, and D7-committed cells and ESCs confirm their close relationship to ESCs (Fig. 3D). By qPCR analysis of select genes, we show that they are indeed exclusively expressed in day 7 cultures (fig. S6C). We also examined several published RNA-seq or microarray datasets of the reprogramming processes and found that these ‘PC candidates trait’ genes are expressed at very low, or undetectable level until the very last reprogramming timepoints, suggesting that these genes could serve as markers for very late stage pluripotency and chimera competency (fig. S6D).

**Fig 3.**
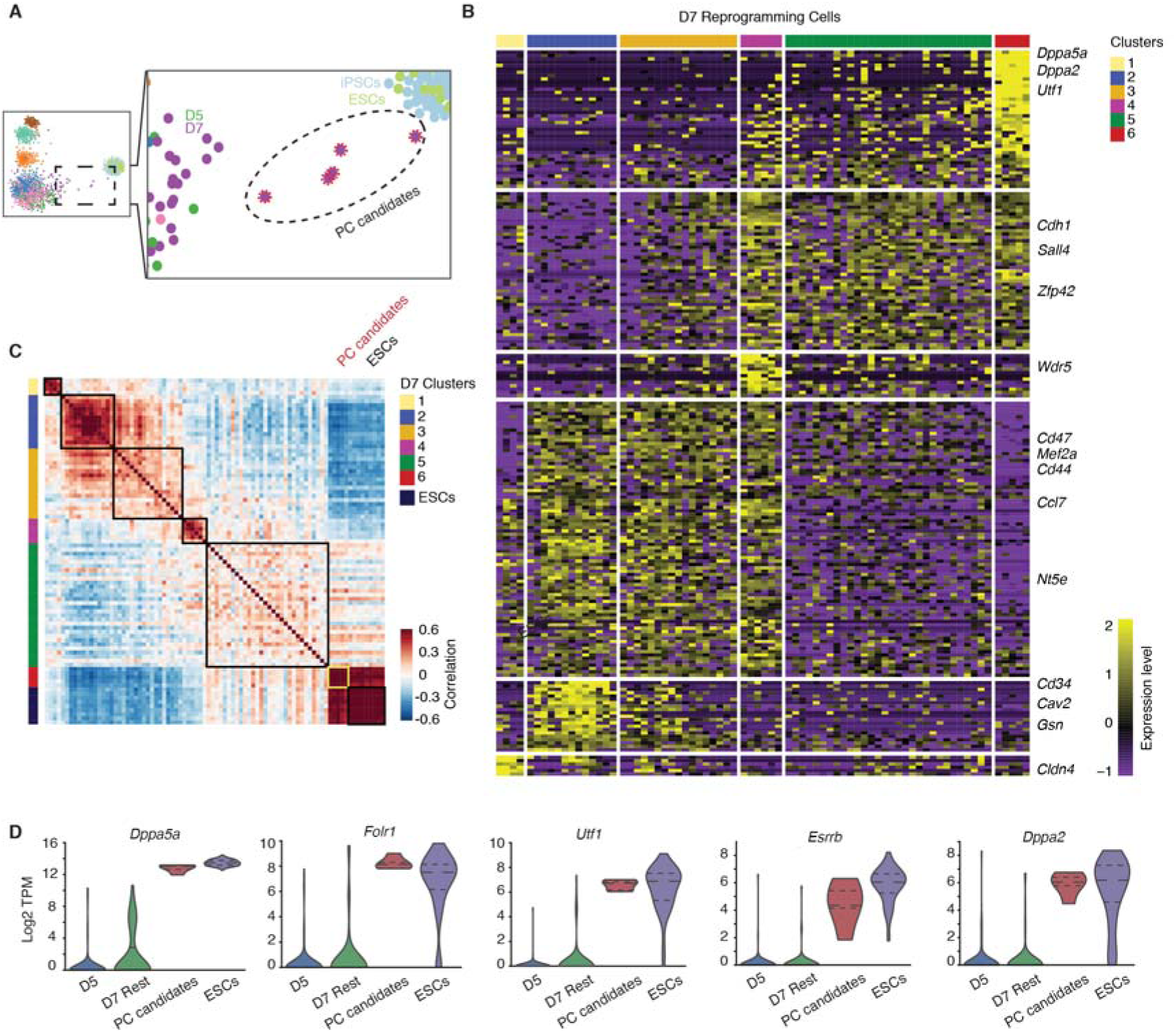
Characterization of the final stages of commitment to pluripotency. **(A)** 5 ‘PC candidates’ as determined from the t-SNE from Fig. 1D reveals 5 D7 cells (in the ellipse) separated from its group and close to ESCs. **(B)** Expression of 210 discriminant genes for 6 sub groups in D7 cells. (**C**) Heatmap shows pair-wise Pearson’s correlation coefficients of 6 sub groups in D7 cells and ESCs. **(D)** The expression pattern of selected PC candidates trait genes at post-infection D5, D7 PC candidates coming from cluster 6 in Fig. 3B, other D7 cells (D7 rest), and ESCs.

### *Dppa5a* as a key marker for chimera competent fate

Based on the heterogeneity of D7 cells, we then attempted to isolate the cluster 6 cells which may represent the final stage of reprogramming and chimera competency for iPSCs. Given its high level of expression in the PC candidates, ESCs, and iPSCs, *Dppa5a* appears to be a strong candidate for a new marker to identify these chimera competent iPSCs. To test this idea, we generated a *Dppa5a*-tdTomato reporter in the OG2 background. We show that colonies at D7 can be identified as *Oct4*-GFP+ and *Dppa5a*-tdTomato+ double positive (G+R+) (Fig. 4A). We further show by FACS analysis of cells at D7 and D8 that *Dppa5a-*tdTomato positive cells can emerge only from the *Oct4*-GFP positive cells (Fig. 4B). We stained the D7 cells with NANOG antibody and showed that *Dppa5a*-tdTomato colonies were emerged from the NANOG positive colonies (fig. S6E). To confirm that G+R+ cells are responsible for the live chimeras, we then performed chimera assay again by picking G+R+ or G+R- colonies at D7.5, and injected them into blastocysts. We showed that the G+R+ cells are capable, but G+R- not, of giving rise to live chimeras (Fig. 4, C and D, and Table S2). Taken together, these results support the idea that the cluster 6 PC candidates defined by single-cell analysis are the ones chimera competent as shown in Figure 1.

**Fig 4.**
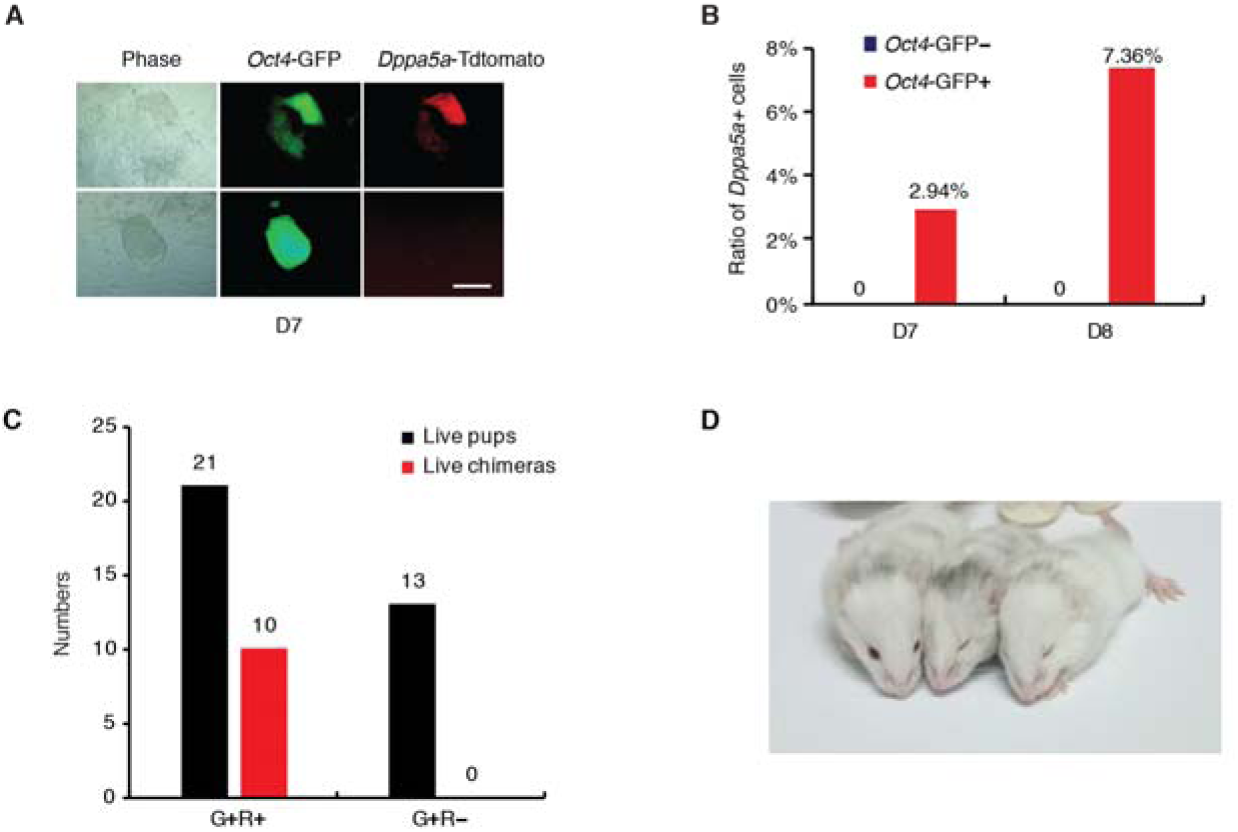
*Dppa5a* as a key marker of later reprogramming stage and chimera competency. **(A)** *Dppa5a*-tdTomato positive colonies emerged from *Oct4*-GFP positive colonies at post-infection D7. Scale bar, 250 µm. **(B)** FACS analysis at post-infection D7 and D8 shows *Dppa5*-tdTomato positive cells emerged from *Oct4*-GFP positive cells. **(C)** Chimera results shows the chimera component iPSCs were *Oct4*-GFP and *Dppa5a*-tdTomato double positive (G+R+) colonies. Data is from four independent experiments (see Table S2). **(D)** Chimera mice derived from *Oct4*-GFP and *Dppa5a*-tdTomato double positive (G+R+) colonies.

### Immune response at the mid to late stage of reprogramming

In addition to discovering late stage markers, we also observed many transient intermediate stages, as has other groups (*15, 34, 40*). These transient stages may be genuine cell state traversals, but may also be barriers to reprogramming. To identify potential barriers preventing the generation of chimera competent iPSCs, we compared specific genes up- or down-regulated between PC candidates vs the rest of the D7 cells (Fig. 5A). Surprisingly the genes most associated with the rest of the D7 cells are related to immune response, and specifically an interferon-gamma response (Fig. 5B). To confirm this, we re-analyzed gene expression from the single cell sequencing, and identified one of the clusters as being primarily immune response genes specifically activated in the middle to late stages of reprogramming (D2 to D7), but silenced at the end stage of reprogramming and in ESCs (Fig. 5, A, C-F, and fig. S7A).

**Fig 5.**
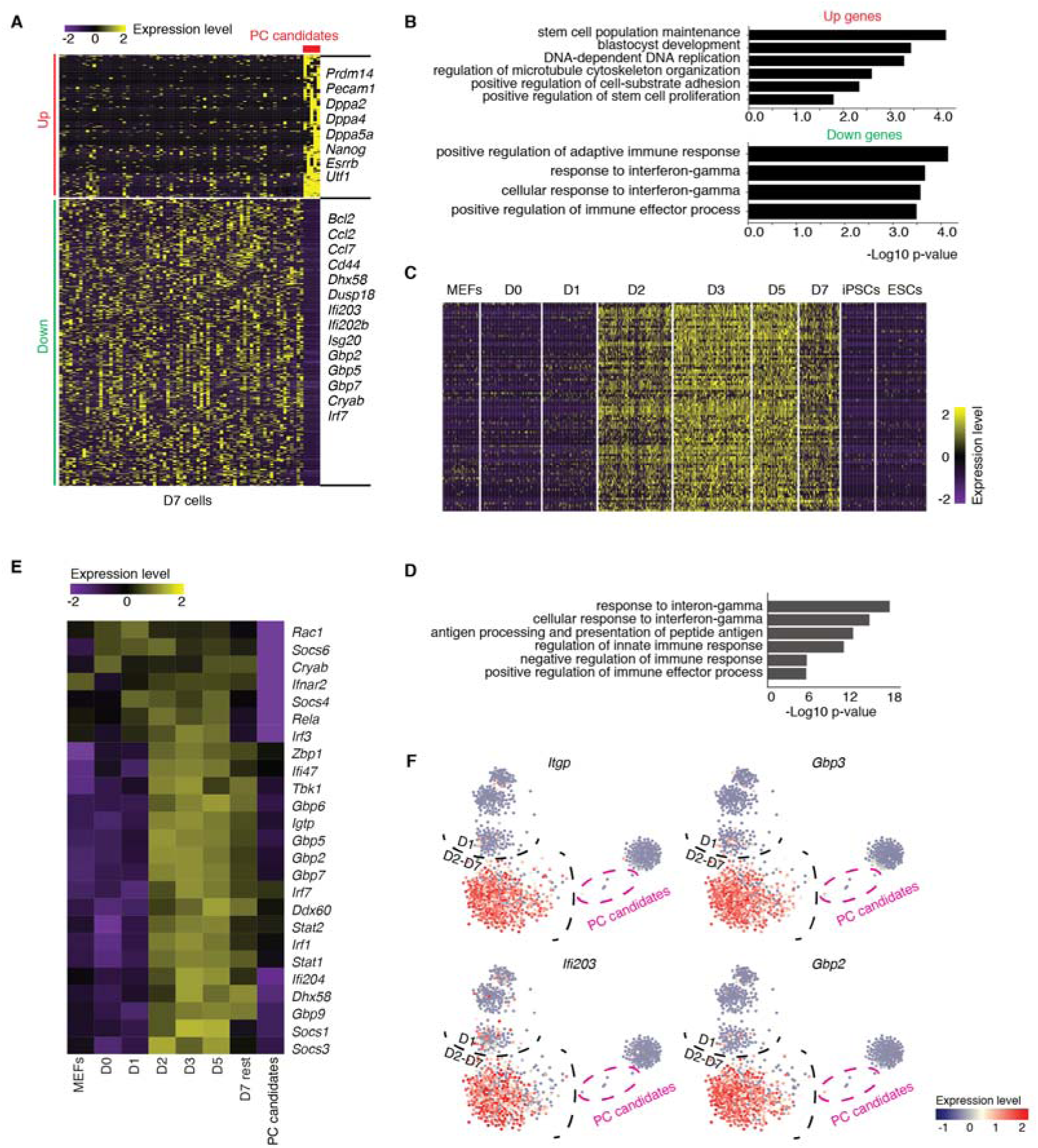
Detection of immune response genes during reprogramming. **(A)** Specific up-regulated and down-regulated genes of PC candidates. Heatmap shows ‘PC candidates’feature genes. Up genes n=182; down genes, n=368, committed cells compared to day 7 rest cells, t-test, p < 1e-5. **(B)** Gene Ontology analysis for genes in Fig. 5A. **(C)** Interferon-gamma response specifically activated at middle stage of reprogramming. Heatmap shows gene expression pattern from cluster marked in red rectangle in fig. S7A, which specific activated in the middle stage during reprogramming. (**D**) Gene Ontology analysis for genes in Fig. 5C. **(E)** Representative immune response genes (IRGs) activated at D2 and silenced in committed cells, heatmap of the selected IRGs, PC candidates are descripted in Fig. 3A. **(E)** t-SNE plot from Fig. 1D, color based on the marker genes expression level.

### IFN-γ impedes the transition to iPSCs

To understand what is causing this immune response we performed ELISA assays on serum free media collected during reprogramming. As shown in Figure 6A, IFN-γ starts to rise at D3 and peaks at D4 and D5, then declines around D7, consistent with the single cell sequencing and qPCR data (Fig. 5, C and D, and fig. S7, A and C). Consistently, we did not observe any appreciable activation of IFN-β (fig. S7B). To see if IFN-γ influences cell fate decision during reprogramming, we added IFN-γ and IFN-β to reprogramming cells. We show that the two cytokines activate different immune response genes (*41*)(Fig. 6B), but critically, only IFN-γ can suppress the expression of late-stage genes, such as *Dppa5a* (Fig. 6C). We further show that IFN-γ completely blocked the formation of *Dppa5a*-Tdtomato positive clones, but only has a limited impact on the formation of *Oct4*-GFP+ colonies (Fig. 6D). Interestingly, we tried to block IFN-γ with anti-IFN-γ antibodies and show that while the antibodies do not enhance *Oct4*-GFP+ colonies, but nevertheless enhance the expression of late reprogramming markers such as *Dppa5a* and *Ooep* (Fig. 6E). However, these genes failed to be up-regulated by antibodies for IFNAR1 (one of the receptor of IFN-β) (fig. S7, D to F), indicating IFN-γ, but not IFN-β, is a barrier for the very late stage of reprogramming we have identified here, marked by *Dppa5a* expression. We also re-analyzed the CD34+ lineage to see if it was responsible for the production of IFN-γ, however, found no such evidence (fig. S7G). These results demonstrate that type-II interferon can act as a barrier to final step of reprogramming.

**Fig 6.**
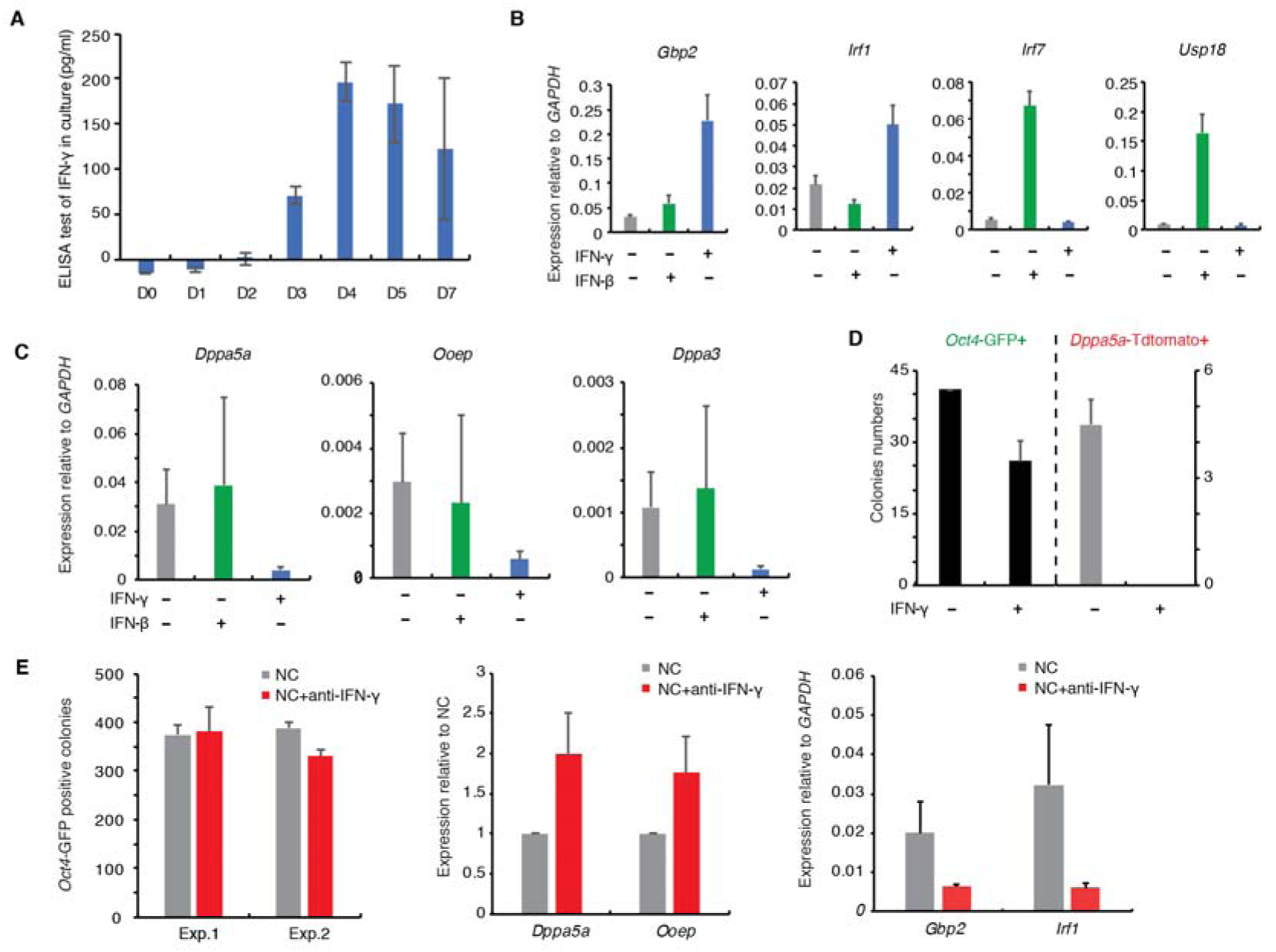
IFN-γ is a barrier for the transition to late reprogramming stage and chimera competency. **(A)** IFN-γ in cell culture supernatant was quantified by ELISA kit, n = 2, biological repeats. **(B) (C)** IFN-γ (20 ng/ml) or IFN-β (20 ng/ml) was added in the medium during generation of iPSCs. qRT-PCR analysis the expression of IFN downstream genes and PC candidates feature genes, n = 3, biological repeats. **(D)** OCT4/SOX2/KLF4 induced OD MEF reprogramming. *Oct4*-GFP and *Dppa5a*-tdTomato positive colonies were counted separately at post-infection day7, with or without the treatment of IFN-γ (20 ng/ml). **(E)** *Oct4*-GFP positive colonies were analyzed after the treatments of IFN-γ antibodies (1 mg/ml). qRT-PCR analysis showed the expression of IFN-γ downstream genes and committed cells feature genes at OCT4/SOX2/KLF4 post-infection D7. n = 2, biological repeats.

## Discussion

In this study, by performing single cell RNA sequencing, we have resolved cell fate transitions during somatic reprogramming. We demonstrate that cells branch into two distinct fates early and CD34 denotes this critical bifurcation to pluripotent or non-pluripotent fates. Towards the pluripotent fate, we identified a late rate-limiting step regulated by IFN-γ and *Dppa5a* as a reliable marker for chimera competent iPSCs. These mechanistic understandings have evaded previous investigation, thus, demonstrating the power of single cell RNA sequencing.

Unlike cells collected in tissue or during development process, cells undergoing somatic reprogramming, which is an artificial process induced by overexpression of transgenes, would render their heterogeneity with more subtle change. As shown in Figure 1D, the reprogramming cells at D2-D7 were scattered and hard to been divided into different cell populations. Another challenge is to confirm the physiology function of cell sub-populations which were defined based on transcriptome data analysis. We set the analyzed results as hypothesizes and proved them experimentally. For example, we defined the “PC candidates” and sorted these cells sub-population with knock-in reporter, then injected them into blastocysts and finally established its chimera-competent function *in vivo*. This strategy assured we got solid understanding about cell heterogeneity during somatic reprogramming.

We and others have argued previously that reprogramming represents one of the best systems to study cell fate conversion mechanistically. Since reprogramming represents cell fate transitions from a fully differentiated cell to a fully pluripotent one, thus, displaying the whole spectrum of possibilities between somatic and pluripotent fates, any mechanism uncovered may become useful and relevant to biology. Our finding that IFN-γ impedes the last transition to pluripotency suggests that stem cell fate and immune responses are interconnected. Indeed, it is interesting to point out that earlier reports have identified components of innate immune response activated by the reprogramming factors or viral infection as a positive event required for the acquisition of pluripotency. For example, toll-like receptor 3 (TLR3) have been shown to be required for reprogramming (*42*), and interleukin-6 (IL-6) can facilitate reprogramming (*43*). It is plausible that innate immunity influences cell fate differentially through individual components such as TLR3, IL6 and IFN-γ. For example, IFN-γ may block the activation of endogenous retroviral elements or other transposable elements so that it prevents the activation of the pluripotent state (*44*). It is expected that more mechanistic insights can be generated that may inform us on how to improve reprogramming technologies.

## Acknowledgments

We appreciated Xiangjie Zhao, Yongqiang Chen, Rongping Luo and Shijuan Huang’s assist on experiment. We are grateful to Lei Gu, Wanqiang Sheng, Yunhao Tan for their helpful discussion and advise. All single cell RNA-sequencing data reported in this study are available at the Gene Expression Omnibus (GEO) under accession GSE103221.

## Supplementary Materials

Materials and Methods

Figures S1-S7

References (*45-50*)

**Other Supplementary Materials for this manuscript includes the following:**

Tables S1 to S9 (Excel format)

## References and Notes

1. K. Takahashi, S. Yamanaka, Induction of pluripotent stem cells from mouse embryonic and adult fibroblast cultures by defined factors. Cell 126, 663–676 (2006).

2. K. Takahashi et al., Induction of pluripotent stem cells from adult human fibroblasts by defined factors. Cell 131, 861–872 (2007).

3. M. Wernig et al., In vitro reprogramming of fibroblasts into a pluripotent ES-cell-like state. Nature 448, 318–324 (2007).

4. K. Takahashi, S. Yamanaka, A decade of transcription factor-mediated reprogramming to pluripotency. Nat Rev Mol Cell Biol 17, 183–193 (2016).

5. J. Yu et al., Induced pluripotent stem cell lines derived from human somatic cells. Science 318, 1917–1920 (2007).

6. M. Nakamura, H. Okano, Cell transplantation therapies for spinal cord injury focusing on induced pluripotent stem cells. Cell Res 23, 70–80 (2013).

7. Y. Avior, I. Sagi, N. Benvenisty, Pluripotent stem cells in disease modelling and drug discovery. Nat Rev Mol Cell Biol 17, 170–182 (2016).

8. B. Xiao, H. H. Ng, R. Takahashi, E. K. Tan, Induced pluripotent stem cells in Parkinson’s disease: scientific and clinical challenges. J Neurol Neurosurg Psychiatry 87, 697–702 (2016).

9. F. Soldner, R. Jaenisch, Medicine. iPSC disease modeling. Science 338, 1155–1156 (2012).

10. I. J. Fox et al., Stem cell therapy. Use of differentiated pluripotent stem cells as replacement therapy for treating disease. Science 345, 1247391 (2014).

11. J. T. Hinson et al., HEART DISEASE. Titin mutations in iPS cells define sarcomere insufficiency as a cause of dilated cardiomyopathy. Science 349, 982–986 (2015).

12. Y. Buganim, D. A. Faddah, R. Jaenisch, Mechanisms and models of somatic cell reprogramming. Nat Rev Genet 14, 427–439 (2013).

13. Z. D. Smith, C. Sindhu, A. Meissner, Molecular features of cellular reprogramming and development. Nat Rev Mol Cell Biol 17, 139–154 (2016).

14. P. Samavarchi-Tehrani et al., Functional genomics reveals a BMP-driven mesenchymal-to-epithelial transition in the initiation of somatic cell reprogramming. Cell Stem Cell 7, 64–77 (2010).

15. Y. Buganim et al., Single-cell expression analyses during cellular reprogramming reveal an early stochastic and a late hierarchic phase. Cell 150, 1209–1222 (2012).

16. J. M. Polo et al., A molecular roadmap of reprogramming somatic cells into iPS cells. Cell 151, 1617–1632 (2012).

17. C. Chronis et al., Cooperative Binding of Transcription Factors Orchestrates Reprogramming. Cell 168, 442–459 e420 (2017).

18. D. Cacchiarelli et al., Integrative Analyses of Human Reprogramming Reveal Dynamic Nature of Induced Pluripotency. Cell 162, 412–424 (2015).

19. R. Li et al., A mesenchymal-to-epithelial transition initiates and is required for the nuclear reprogramming of mouse fibroblasts. Cell Stem Cell 7, 51–63 (2010).

20. A. Soufi, G. Donahue, K. S. Zaret, Facilitators and impediments of the pluripotency reprogramming factors’ initial engagement with the genome. Cell 151, 994–1004 (2012).

21. P. D. Tonge et al., Divergent reprogramming routes lead to alternative stem-cell states. Nature 516, 192–197 (2014).

22. D. Cacchiarelli et al., Integrative Analyses of Human Reprogramming Reveal Dynamic Nature of Induced Pluripotency. Cell 162, 412–424 (2015).

23. B. Di Stefano et al., C/EBPalpha creates elite cells for iPSC reprogramming by upregulating Klf4 and increasing the levels of Lsd1 and Brd4. Nat Cell Biol 18, 371–381 (2016).

24. J. Chen et al., H3K9 methylation is a barrier during somatic cell reprogramming into iPSCs. Nat Genet 45, 34–42 (2013).

25. J. Chen et al., Vitamin C modulates TET1 function during somatic cell reprogramming. Nat Genet 45, 1504–1509 (2013).

26. B. Papp, K. Plath, Epigenetics of reprogramming to induced pluripotency. Cell 152, 1324–1343 (2013).

27. T. W. Theunissen, R. Jaenisch, Molecular control of induced pluripotency. Cell Stem Cell 14, 720–734 (2014).

28. K. Hochedlinger, R. Jaenisch, Induced Pluripotency and Epigenetic Reprogramming. Cold Spring Harb Perspect Biol 7, (2015).

29. D. Grun, A. van Oudenaarden, Design and Analysis of Single-Cell Sequencing Experiments. Cell 163, 799–810 (2015).

30. C. Gawad, W. Koh, S. R. Quake, Single-cell genome sequencing: current state of the science. Nat Rev Genet 17, 175–188 (2016).

31. A. Tanay, A. Regev, Scaling single-cell genomics from phenomenology to mechanism. Nature 541, 331–338 (2017).

32. L. Wen, F. Tang, Single-cell sequencing in stem cell biology. Genome Biol 17, 71 (2016).

33. J. Chen et al., Rational optimization of reprogramming culture conditions for the generation of induced pluripotent stem cells with ultra-high efficiency and fast kinetics. Cell Res 21, 884–894 (2011).

34. E. Lujan et al., Early reprogramming regulators identified by prospective isolation and mass cytometry. Nature 521, 352–356 (2015).

35. J. Fan et al., Characterizing transcriptional heterogeneity through pathway and gene set overdispersion analysis. Nat Methods 13, 241–244 (2016).

36. L. Haghverdi, M. Buttner, F. A. Wolf, F. Buettner, F. J. Theis, Diffusion pseudotime robustly reconstructs lineage branching. Nat Methods 13, 845–848 (2016).

37. X. Qiu et al., Single-cell mRNA quantification and differential analysis with Census. Nat Methods 14, 309–315 (2017).

38. X. Qiu et al., Reversed graph embedding resolves complex single-cell developmental trajectories. bioRxiv, (2017).

39. C. Xu, Z. C. Su, Identification of cell types from single-cell transcriptomes using a novel clustering method. Bioinformatics 31, 1974–1980 (2015).

40. E. R. Zunder, E. Lujan, Y. Goltsev, M. Wernig, G. P. Nolan, A continuous molecular roadmap to iPSC reprogramming through progression analysis of single-cell mass cytometry. Cell Stem Cell 16, 323–337 (2015).

41. W. M. Schneider, M. D. Chevillotte, C. M. Rice, Interferon-stimulated genes: a complex web of host defenses. Annu Rev Immunol 32, 513–545 (2014).

42. J. Lee et al., Activation of innate immunity is required for efficient nuclear reprogramming. Cell 151, 547–558 (2012).

43. L. Mosteiro et al., Tissue damage and senescence provide critical signals for cellular reprogramming in vivo. Science 354, (2016).

44. A. P. Hutchins, D. Pei, Transposable elements at the center of the crossroads between embryogenesis, embryonic stem cells, reprogramming, and long non-coding RNAs. Sci Bull (Beijing) 60, 1722–1733 (2015).

